# High-throughput profiling and analysis of plant responses over time to abiotic stress

**DOI:** 10.1101/132787

**Authors:** Kira M. Veley, Jeffrey C. Berry, Sarah J. Fentress, Daniel P. Schachtman, Ivan Baxter, Rebecca Bart

## Abstract

Sorghum (*Sorghum bicolor* (L.) Moench) is a rapidly growing, high-biomass crop prized for abiotic stress tolerance. However, measuring genotype-by-environment (G × E) interactions remains a progress bottleneck. Here we describe strategies for identifying shape, color and ionomic indicators of plant nitrogen use efficiency. We subjected a panel of 30 genetically diverse sorghum genotypes to a spectrum of nitrogen deprivation and measured responses using high-throughput phenotyping technology followed by ionomic profiling. Responses were quantified using shape (16 measurable outputs), color (hue and intensity) and ionome (18 elements). We measured the speed at which specific genotypes respond to environmental conditions, both in terms of biomass and color changes, and identified individual genotypes that perform most favorably. With this analysis we present a novel approach to quantifying color-based stress indicators over time. Additionally, ionomic profiling was conducted as an independent, low cost and high throughput option for characterizing G × E, identifying the elements most affected by either genotype or treatment and suggesting signaling that occurs in response to the environment. This entire dataset and associated scripts are made available through an open access, user-friendly, web-based interface. In summary, this work provides analysis tools for visualizing and quantifying plant abiotic stress responses over time. These methods can be deployed as a time-efficient method of dissecting the genetic mechanisms used by sorghum to respond to the environment to accelerate crop improvement.

## INTRODUCTION

The selection of efficient, stress-tolerant plants is essential for tackling the challenges of food security and climate change, particularly in hot, semiarid regions that are vulnerable to economic and environmental pressures (Lobell et al., 2008; Foley et al., 2011; DeLucia et al., 2014; Hadebe et al., 2016). Many crop species, having undergone both natural and human selection, harbor abundant, untapped genetic diversity. This genetic diversity will be a valuable resource for selecting and breeding crops to maximize yield under adverse environmental conditions (Leakey, 2009). Sorghum (*Sorghum bicolor* (L.) Moench) originated in northern Africa and was domesticated 8,000 – 10,000 years ago. Thousands of genotypes displaying a wide range of phenotypes have been collected and described (Deu et al., 2006; Paterson et al., 2009; Lasky et al., 2015). *Sorghum bicolor*, the primary species in cultivation today, has many desirable qualities including the ability to thrive in arid soils with minimal inputs, and many end-uses (Morris et al., 2013; Vermerris and Saballos, 2013). For example, grain varieties are typically used for food and animal feed production, sweet sorghum genotypes accumulate non-structural, soluble sugar for use as syrup or fuel production, and bioenergy sorghum produces large quantities of structural, lignocellulosic biomass that may be valuable for fuel production (Murray, 2013; Rooney, 2014). Sorghum genotypes can be differentiated and categorized by type according to these end-uses.

Rising interest in sorghum over the last forty years has led to efforts to preserve and curate its diversity. To maximize utility, these germplasm collections must now be characterized for performance across diverse environments (Furbank and Tester, 2011; Fiorani and Schurr, 2013; Araus and Cairns, 2014). Deficits in our understanding of genotype-by-environment interactions (G × E = P, where G = genotype, E = environment and P = phenotype) are limiting current breeding efforts (Zamir, 2013). Controlled-environment studies are quantitatively robust but are often viewed with skepticism regarding their translatability to field settings. Further, they can often accommodate only a limited number of genotypes at a time. In contrast, field level studies allow for large numbers of genotypes to be evaluated simultaneously. However, these studies provide limited resolution to resolve the effect of environment on phenotype and often require multi-year replication. This conundrum has motivated enthusiasm for both controlled environment and field level high throughput phenotyping platforms. However, the use of large-scale phenotyping and statistical modeling to predict field-based outcomes is challenging (Deans et al., 2015; Lipka et al., 2015; Zivy et al., 2015).

Here, we sought to define a set of measurable, environmentally-dependent, phenotypic outputs to aid crop improvement. We utilized automated phenotyping techniques under controlled-environmental conditions to characterize G × E interactions on a diverse panel of sorghum genotypes in response to abiotic stress. Specifically, we describe and quantify statistically robust differences among the genotypes to nutrient-poor conditions using three phenotypic characteristics: biomass, color, and ion accumulation. Using image analysis to characterize leaf color and biomass over time in conjunction with ionomics, we report measurable, genetically-encoded, phenotypic traits that are affected by nitrogen treatment. This work presents a foundation for understanding the range of sorghum early-responses to abiotic stress and provides tools for analyzing other available datasets.

## RESULTS

### Phenotypic effects of nitrogen treatment on a sorghum diversity panel

Next to water, nutrient supply (most notably nitrogen availability) is often cited as the most important environmental factor constraining plant productivity (Chapin et al., 1987; Liu et al., 2015). The initial goal of our experimental design was to enable the early detection and quantification of stress responses in plants. Figure 1 illustrates the overall experimental design we used to test the phenotypic effects of nitrogen treatment on sorghum over the course of a three-week-long experiment using high-throughput phenotyping. Three nitrogen treatments were designed to analyze the effects of source (i.e. ammonium vs. nitrate) and quantity of nitrogen on plant development over time (Figure 1A, methods). For this study, sorghum was chosen for its genetic diversity and wide range of abilities to thrive under semi-arid, nutrient-limited conditions. In order to test the role that genotype plays in response to nitrogen treatment, a panel of 30 sorghum lines was assembled (Table S1). This panel includes sorghum accessions from all five cultivated races (bicolor, caudatum, durra, guinea and kafir), representing a variety of geographic origins and morphologies (Kimber et al., 2013; Brenton et al., 2016). The genotypes also display a range of photoperiod sensitivities and are categorized into three general production types: grain, sweet, and bioenergy. This diversity was intended to generate a range of responses that could be measured and attributed to either genotype, stress treatment, or both.

**Figure 1.**
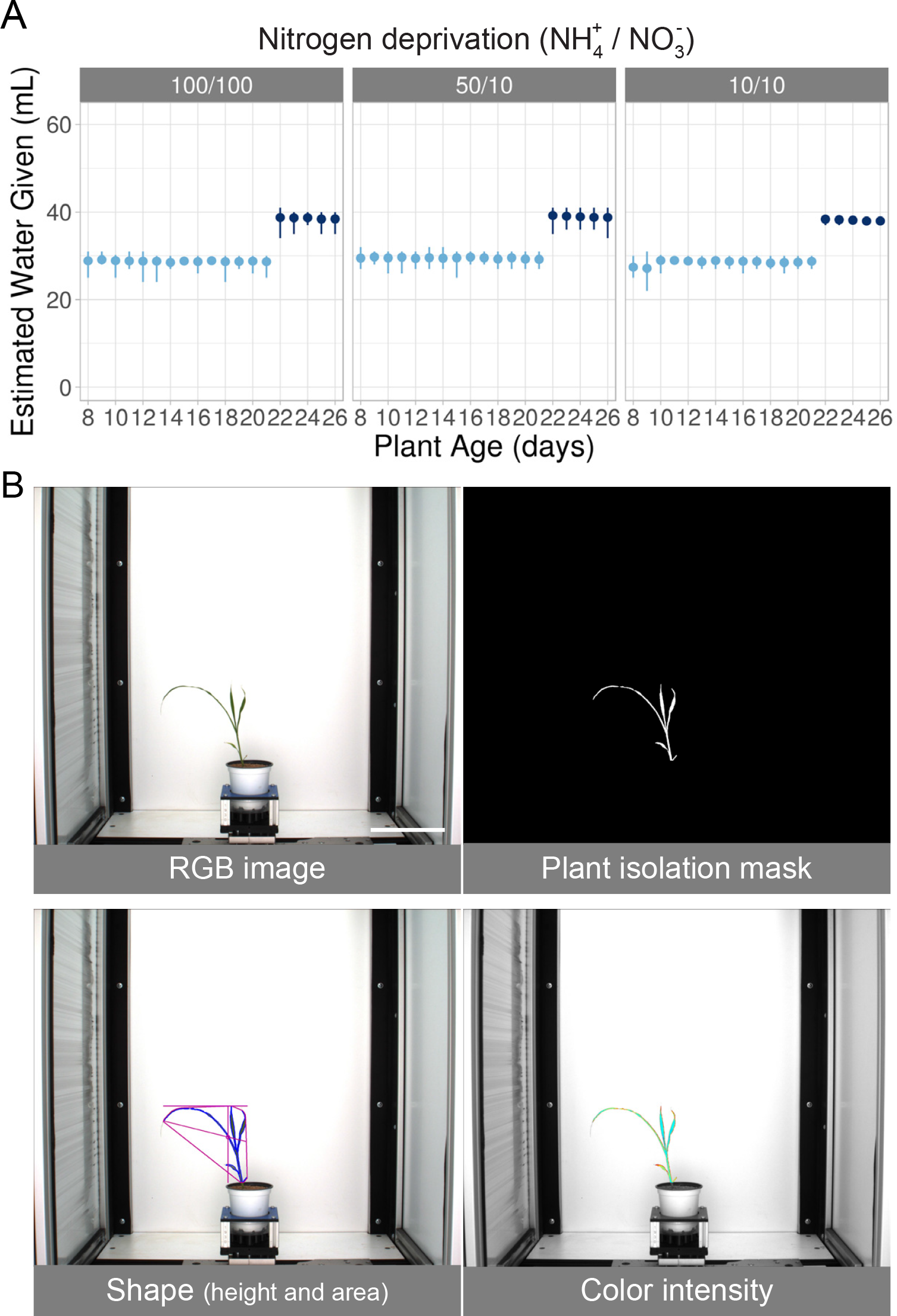
Experimental Overview. A) Watering regime used for nitrogen deprivation. The x-axis shows the age of the plants throughout the experiment and the y-axis indicates the estimated volume of water plus nutrients (ml), calculated based on the weight change of the pot before and after watering. Each dot represents the average amount of water delivered each day with vertical lines indicating error (99% confidence interval). Watering regime was increased due to plant age (shades of blue). The experimental treatments are listed above the plots. Volume of water and source of nitrogen are indicated and was scaled based on the 100% (100/100) treatment group (1 mM ammonium / 14.5 mM nitrate for 100% treatment group). B) Image analysis example (genotype NTJ2 from 100/100 treatment group on day 16 is shown). Top row: Example original RGB image taken from phenotyping system and plant isolation mask generated using PlantCV. Bottom row: two examples of attributes analyzed (area and color). Scale bar = 15 cm.

With the use of automated phenotyping, all plants were photographed daily and images were processed using the open source PlantCV analysis software package ((Fahlgren et al., 2015), http://plantcv.danforthcenter.org). Within each RGB image, the plant material was isolated, allowing phenotypic attributes to be analyzed (Figure 1B). Scripts used to make the figures within this manuscript, along with the raw data, are available here: http://plantcv.danforthcenter.org/pages/data-sets/sorghum_abiotic_stress.html. In total, 16 different shape characteristics were quantified (Figure S1). Principal component analysis (PCA) of all the quantified attributes revealed that shape characteristics could be used to separate all three treatments (Figure 2A). Our results indicated that “area” was the plant shape feature that displayed the largest treatment effect. We consider area measurements from plant images as a proxy for biomass measurements as these traits have been shown to be correlated for a number of plant species, including sorghum (Fahlgren et al., 2015; Neilson et al., 2015). Additionally, the effect of low nitrogen on plant color is well established and RGB image-based methods have been described to estimate chlorophyll content of leaves (Hu et al., 2010; Shibghatallah et al., 2013; Wang et al., 2014; Cendrero-Mateo et al., 2016; Junker and Ensminger, 2016; Mishra et al., 2016). In contrast to shape, PCA of color (hue and intensity) attributes at the end of the experiment only separated the high nitrogen treatment group away from the two lower nitrogen treatment groups (Figure 2B). These data indicate that the different nitrate concentrations in the two lower nitrogen treatment groups significantly affects shape but not color. To further explore the effect that our experimental treatments had on the measured shape characteristics and color for each individual genotype, an interactive version of the generated data is available here: (http://plantcv.danforthcenter.org/pages/data-sets/sorghum_abiotic_stress.html).

**Figure 2.**
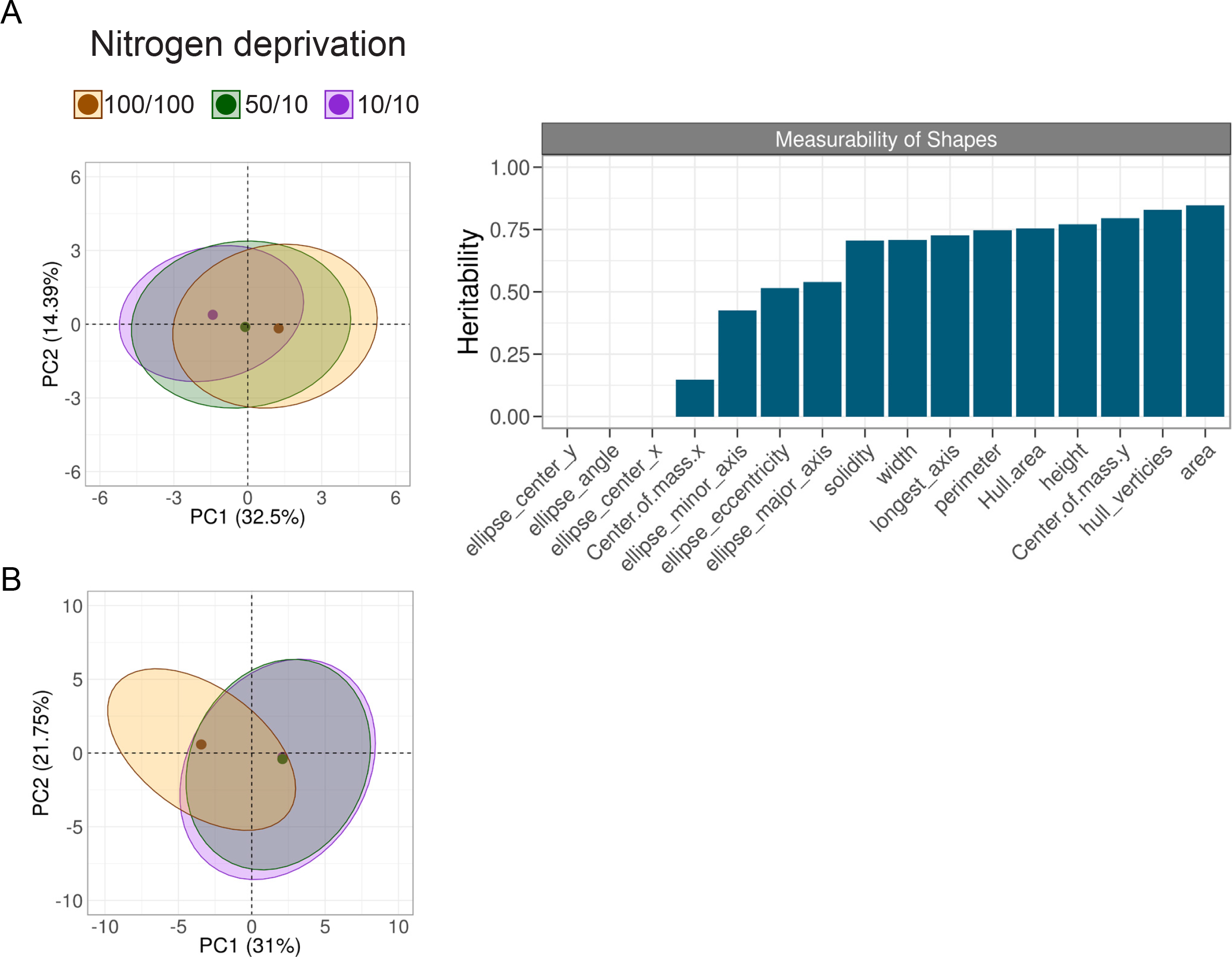
Determining plant attributes affected by experimental treatments. A) Left: Principle Component Analysis (PCA) plots of shape attributes for plants subjected to nitrogen deprivation at the end of the experiment (plant age 26 days). 95% confidence ellipses are calculated for each of the treatment groups and the dots indicate the center of mass. The shape attributes included in the PCA are as follows: area, hull area, solidity, perimeter, width, height, longest axis, center of mass x-axis, center of mass y-axis, hull vertices, ellipse center x-axis, ellipse center y-axis, ellipse major axis, ellipse minor axis, ellipse angle and ellipse eccentricity. Right: Bar graph indicating measurability of shape attributes, showing the proportion of variance explained by treatment (i. e. treatment effect, y-axis). B) PCA plots showing analysis of color values within the mask for plants subjected to nitrogen deprivation at the end of the experiment (plant age 26 days). All 360 degrees of the color wheel were included, binned every 2 degrees

Many factors contribute to the ability of plants to utilize nutrients and presumably, much of this is genetically explained. Correspondingly, genotype was a highly significant variable (*p*-value = 0.003 when measuring area) within this dataset. To investigate how much nitrogen treatment response is explained by major genotypic groupings, we calculated the contribution of type, photoperiod, or race on treatment effect. Of these, photoperiod was the only grouping that significantly contributed to area (Figure S2).

### Size and growth rate during nitrogen stress conditions

Nitrogen stress tolerance is a plant’s ability to thrive in low nitrogen conditions. To identify sorghum varieties tolerant to growth in nutrient limited conditions, we considered plant size at the end of the experiment within the most severe nitrogen deprivation treatment group for all genotypes (Figure 3A). In this experiment, San Chi San, PI_510757, PI_195754, BTx623 and PI_508366 were larger than average as compared to all other genotypes under low nitrogen conditions. In contrast, Della, PI_297155 and PI_152730 were smaller than average. Next we aimed to leverage the temporal resolution available from high throughput phenotyping platforms. For these experiments we considered average growth rate across the experiment (Figure 3B). Overall, end plant size correlated well with overall growth rates. For example, by both measures, Della displayed particularly weak growth characteristics under low nitrogen conditions while BTx623 performed well. However, the correlation was imperfect. San Chi San displayed the largest end size but was statistically average in terms of growth rate across the experiment. Discrepancies between end-biomass and growth rate (e.g. large plants with average or low observed growth rates) may indicate differences in germination rates (e.g. being larger at the beginning of the phenotyping experiment). Taken together, these data suggest that PI_195754, BTx623 and PI_508366 are the best performing genotypes tested under low nitrogen conditions.

**Figure 3.**
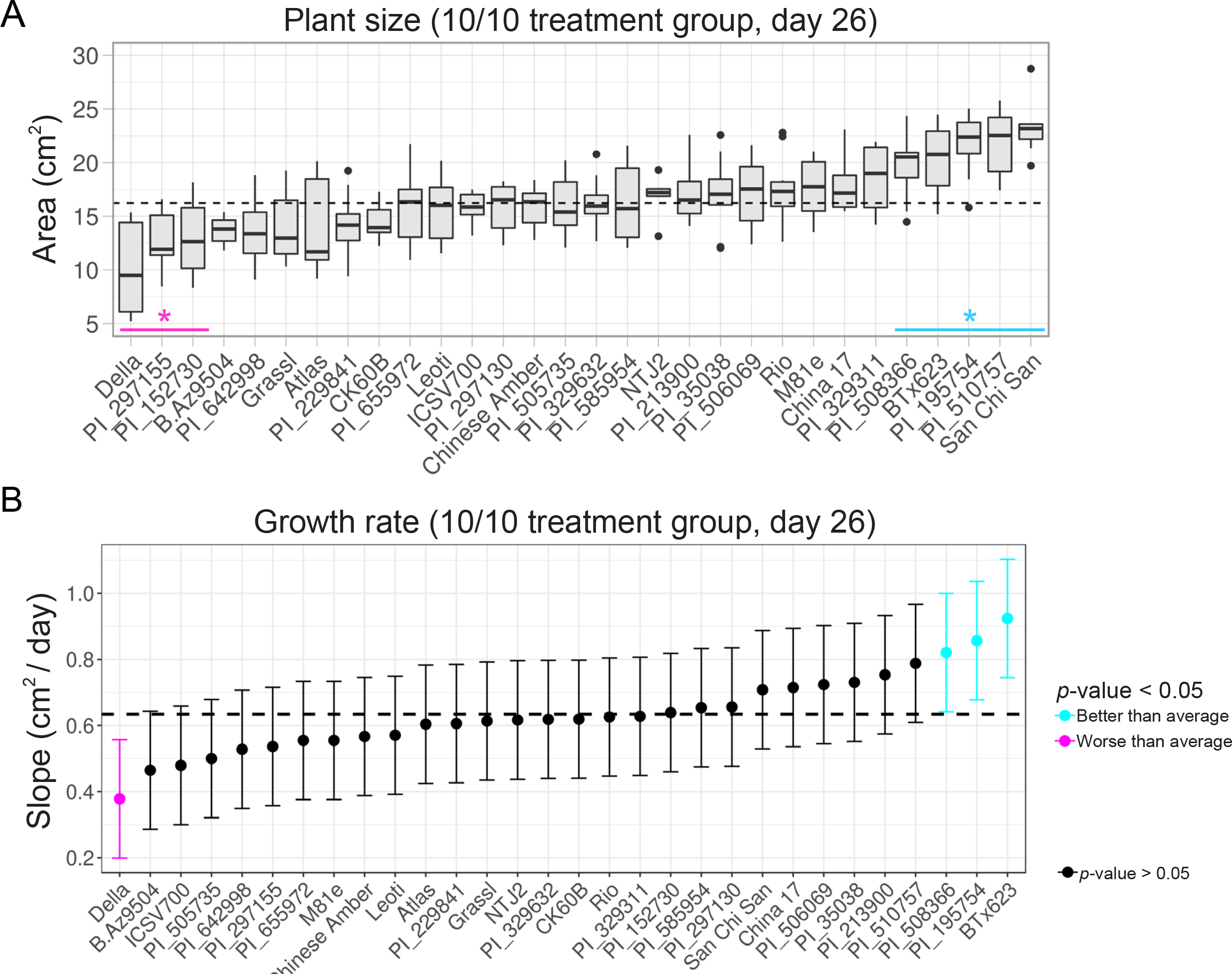
Growth response of genotypes to nitrogen deprivation. A) Boxplot showing average plant size (area) at the end of the experiment (day 26), * q-values < 0.01) with outliers (dots) at the end of the experiment for the 10/10 treatment group. The median is indicated by a black bar within each box. B) Growth rate (average change in area per day, days 10-22) for the 10/10 treatment group. The dotted lines indicate the treatment group average in both panels. Genotypes that displayed greater than average (blue) or less than average (magenta) growth are indicated. Error bars: 95% confidence intervals for both graphs

In contrast to nitrogen stress tolerance, nitrogen use efficiency is often defined as a plant’s ability to translate available nitrogen into biomass. China 17 and San Chi San are considered nitrogen-use-efficient genotypes, while BTx623 and CK60B have previously been reported as less efficient (Maranville and Madhavan, 2002; Gelli et al., 2014, 2017). To further explore nitrogen use efficiency phenotypes within our experiment, we factored timing of growth response differences into our analysis. For each day, we analyzed biomass for each genotype within the 100% control group (100 NH_4_^+^/100 NO_3_^−^) and compared that to the biomass within the 10% treatment group (10 NH_4_^+^/10 NO_3_^−^). Comparing these two populations allowed us to determine when, during the course of our experiment, those figures became significantly different (Figure 4A).

**Figure 4.**
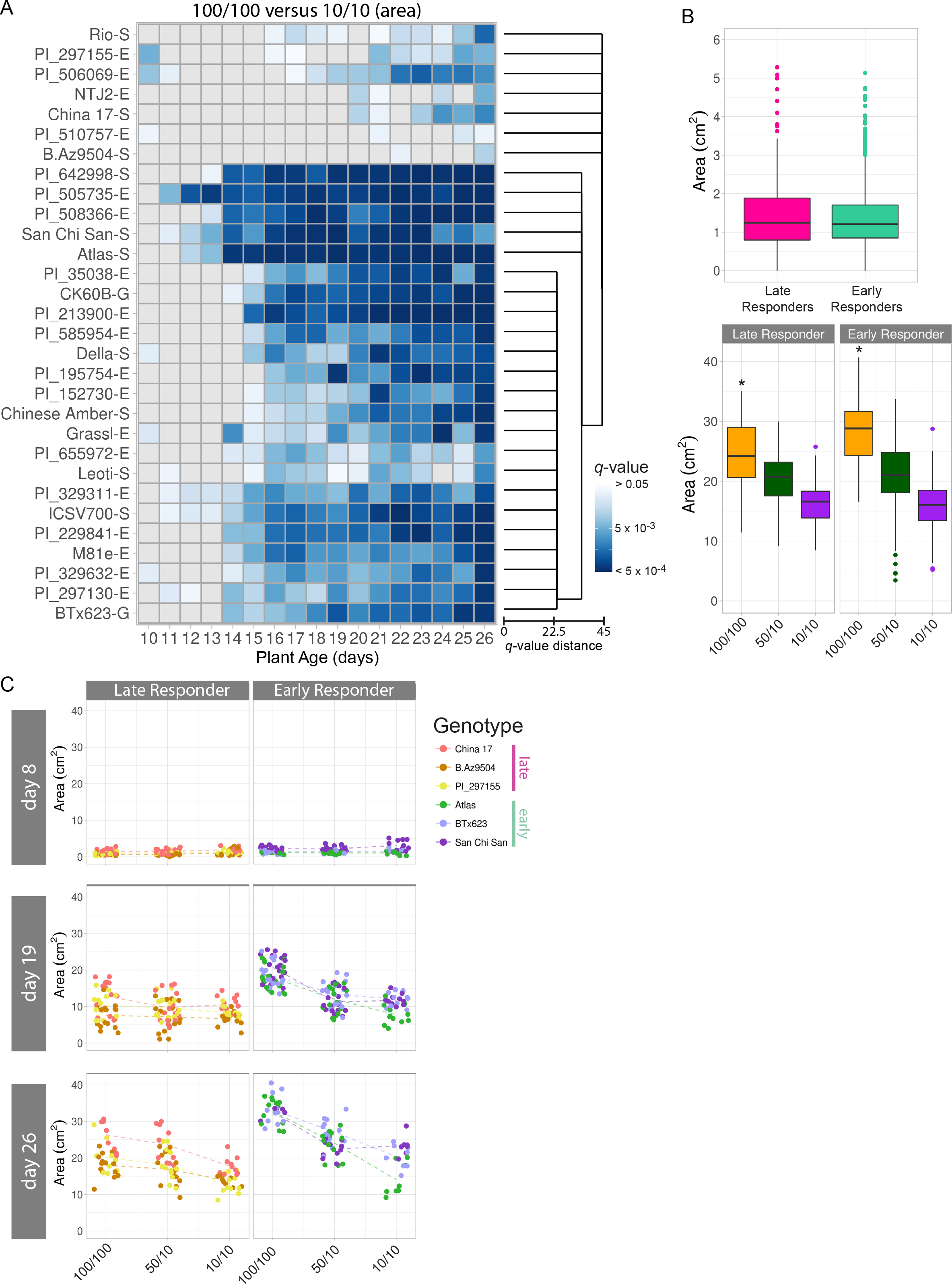
Timing of response to nitrogen: size changes in late and early responding genotypes. A) Statistical analysis of differences in area over time (bottom, plant age) for the 30 sorghum genotypes analyzed. *q*-values for the heat map are indicated in blue, with darkest coloring representing most significance. The Canberra distance-based cluster dendrogram (right) was generated from calculated *q*-values. B) Box plots showing average biomass (area) with outliers (colored dots) for late- (left) and early- (right) responding lines from panel A at the beginning (day 8, top) and end (day 26, bottom) of the experiment. The median is indicated by a black bar within each box. * indicates significant difference between early and late groups (*p*-value < 5 × 10^−6^). C) Scatter plots representing plant area (y-axis) by treatment (x-axis) at the beginning (day 8), middle (day 19), and end (day 26) of the experiment for chosen late responding (left) and early responding (right) genotypes (key, right). Each dot represents an individual plant on a day and dotted lines connect genotypic averages

This analysis separated the genotypes into two broad categories: “early” responding accessions and “late” responding accessions. Early- and late-responding lines were not found to be significantly different in terms of size before treatment administration (Figure 4B, top panel). Therefore, we hypothesized that either 1) lines would be late-responding because they were proficient at using any level of available nitrogen or 2) because they grew slowly regardless of quantity of nitrogen supplied. We found that the early-responding lines were larger, on average, than the late-responding lines within the 100/100 treatment group (Figure 4B, bottom panel) suggesting that these lines are more competent at using available nitrogen. A subset of these genotypes are displayed in Figure 4C to illustrate our observations. The genotype Atlas is an example of a very early responding line, and it was one of the largest plants in the 100/100 treatment group, but also one of the worst-performing lines in the 10/10 treatment group (Figures 3A, 3B, 4C). In contrast, China 17 performed relatively well under nitrogen-limited conditions (10/10), but when nitrogen was abundant (100/100) the biomass accumulation was relatively poor (Figure 3A, 3B, 4C). A similar phenotype was observed for PI_510757. In addition to varying the amount of nitrogen available, we also tested whether any lines harbor a preference for nitrogen source. Nitrogen is typically available in two ionic forms within the soil, ammonium and nitrate, both of which are actively taken up into plant roots by transporters located in the plasma membrane (Crawford and Forde, 2002; Kiba and Krapp, 2016). Expression of these gene products and others have been shown to be responsive to nitrogen availability in sorghum (Vidal et al., 2014). For example, San Chi San and China 17 are known to have higher levels of expression of nitrate transporters when compared to nitrogen-use-inefficient lines (Gelli et al., 2014). Notably, Atlas translated an increased ammonium level into larger plant size. In contrast, San Chi San showed no change in average plant size between the two lower nitrogen treatments (Figure 4C). Among the 30 tested genotypes, 16 displayed little difference between the 50/10 and 10/10 groups in terms of plant size toward the end of the experiment (Figure S3). This highlights the importance of considering both quantity and source when investigating nitrogen responses.

### Combined size and color analysis over time

In addition to affecting shape attributes, nitrogen starvation generally results in reduced chlorophyll content and increased chlorophyll catabolism. Other groups have used image analysis to estimate chlorophyll content and nitrogen use in rice (Wang et al., 2014). The RGB images contain plant hue channel information, and this was found to be a separable characteristic within the nitrogen deprivation treatment groups (Figure 2B). We assessed color-based responses to nitrogen treatment in the early- and late-responding genotypes as defined in Figure 4A (Figure 5). To facilitate this analysis we used the generated histograms of images of the individual plants from each day of the experiment and averaged those from the early and late categories within each treatment group (Figure 5A, day 13). We found that the histograms of the plant images contained two primary peaks: yellow and green. For both early-and late-responding lines, the yellow peak was larger than the green for the plants in the 10/10 treatment group as compared to the 100/100 treatment group. Early-responding lines within the 100/100 treatment group displayed the largest green-channel values. Late responding lines grown under nitrogen-limiting conditions displayed the largest yellow channel values. In order to further visualize color-based treatment effects, we subtracted the 10/10 histograms from the 100/100 histograms and plotted this difference (Figure S4). This revealed that although the late responding lines were more yellow, the magnitude difference from the treatment was similar for early and late lines in the yellow channel. In contrast, the early-responding lines tended to have a larger green channel difference between the 10/10 and the 100/100 treatment groups, with early-responding lines showing a larger difference in the green channel.

**Figure 5.**
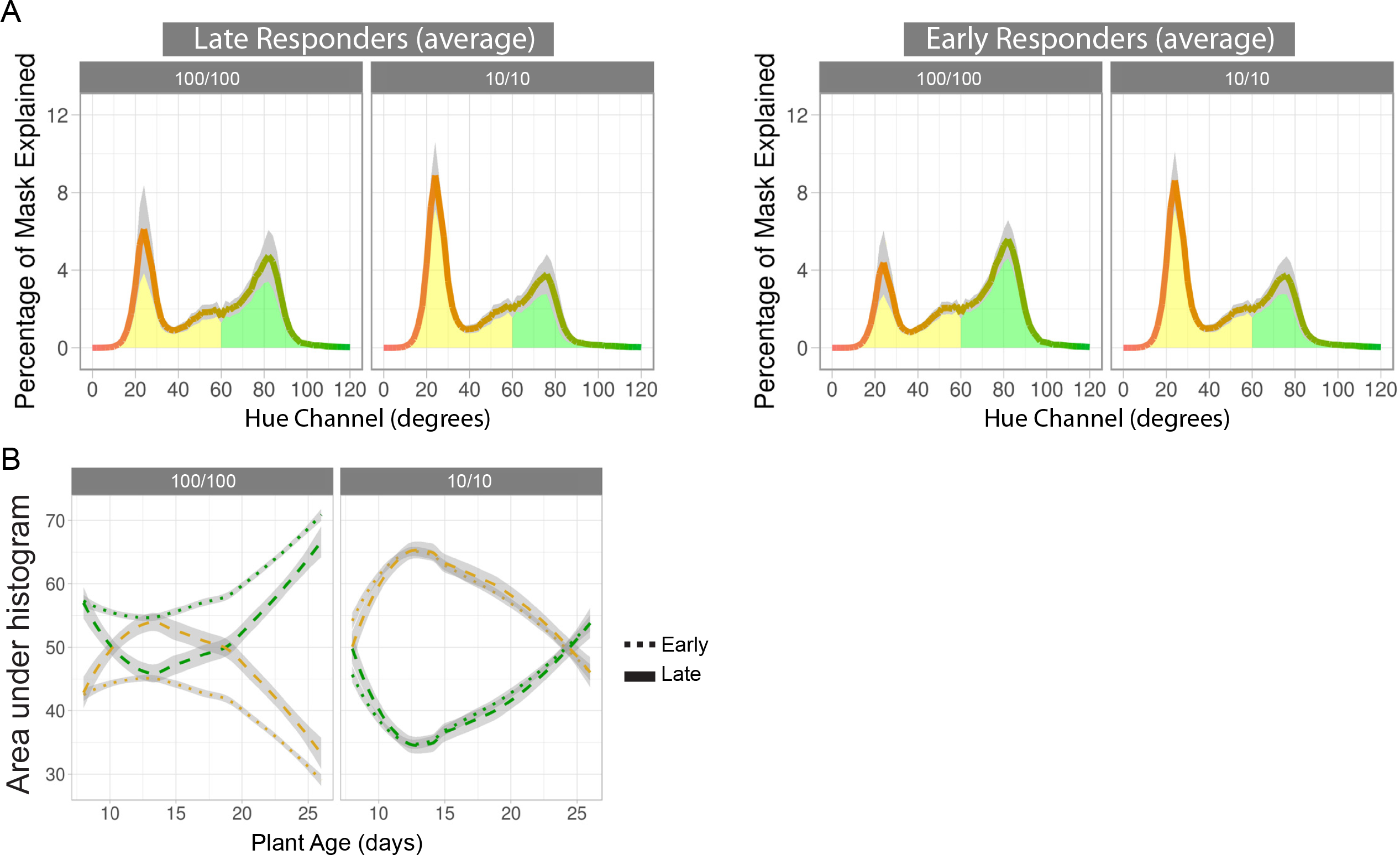
Color changes in late and early responding genotypes to nitrogen treatment. A) Average histograms illustrating percentage of identified plant image mask (y-axis) represented by a particular hue degree (x-axis). Presented is the average of the early- and late-responding lines on day 13 of the experiment. Yellow and green areas of the hue spectrum are highlighted as such. B) Change in yellow (degrees 0 - 60) and green (degrees 61 - 120) hues over time for 100/100 (left) and 10/10 (right) treatment groups. Plotted is the area under the curves presented in A (y-axis) over the duration of the experiment (x-axis) for early- and late-responding genotypes. Grey areas indicate standard error

To assess color-based treatment effects over time, we took the area under the histograms (e. g. Figure 5A) for all time points and plotted them against plant age (Figure 5B). Given the peaks within the histograms mentioned above, we focused on these regions and defined yellow (degrees 0 - 60) and green (degrees 61 - 120) to facilitate quantitative analysis. As expected, plants within the 10/10 treatment group were generally more yellow (and consequently less green) over the course of the stress treatment. We detect a peak difference between yellow and green occurring on day 13, then the effect diminishes. A similar peak and overall pattern is seen in the 100/100 treatment group, with plants greening after day 13. Focusing on either the green or the yellow hue, there was no discernable difference in color over time between late- and early-responding lines within the 10/10 treatment group (left panel, dotted versus dashed lines, *p* > 0.05). However, within the 100/100 treatment group, early-responding lines were consistently greener, while late-responding genotypes became increasingly yellow until day 13. Late- and early-responding lines behaved differently under the 100/100 nitrogen treatment conditions, becoming significantly different quickly (day 10, *p* < 0.05) and remaining so for the duration of the experiment, with the most significant difference occurring on day 13 (*p* < 1 × 10^−15^).

Combining the above plant size- and color-based data, we conclude that the ‘early responding phenotype’ indicates that these plants are able to take better advantage of available nutrients. Importantly, both size and color phenotypes indicate that the early responding genotypes do not display the fitness advantage in low nitrogen conditions. Together, these data demonstrate that color-based image analysis is consistent with and complimentary to the more-established biomass measures of fitness and performance.

### Ionomic profiling as a heritable, independent, measurable readout of abiotic stress

In addition to the image-based analysis used above to reveal measurable size- and color-based outcomes in response to nitrogen treatment, we also performed ionomic analysis to gain better insight into the physiological changes that occur in response to nitrogen (Figure S5). It has been established that both genetic and environmental factors and their interactions play a significant role in determining the plant ionome (Baxter et al., 2008; Baxter and Dilkes, 2012; Chao et al., 2012; Asaro et al., 2016; Shakoor et al., 2016; Thomas et al., 2016). Thus, this analysis was used to explore alterations that might not be revealed by shape or color analysis but would still contribute to the effect of nutrient availability. Each element was modeled as a function of both genotype and treatment, and genotype was a significant factor for most elements with Mo, Cd, and Co being the most affected by genotype (Figures 6, S5) indicating that concentrations of these elements may be the most directly affected by genetically encoded traits. Nitrogen deprivation had a measurable effect on every element (Figure 6A, B). As was seen for color (Figure 2B), PCA of the elements revealed separation of the nitrogen treatments, with the two lower nitrogen treatment groups separating from the high treatment group (Figure 6B). Both micro (Se, Rb, Mo, Cd) and macro (K, P) nutrients contributed strongly to the PCs separating the 100/100 treatment group away from the other two treatments within the PCA. Interestingly, under our experimental conditions, phosphorous was one of the elements with the largest nitrogen treatment effect (Figure 6A). In Arabidopsis, the presence of nitrate has been shown to inhibit phosphorous uptake (Kant et al., 2011; Lin et al., 2013). Consistent with this, dry weight-based concentrations of phosphorous were inversely proportional to administered nitrate treatment, with the 100/100 treatment group accumulating less on average than either 50/10 or 10/10 (Figure 6C, *p* < 1 × 10^−16^, Student’s *t*-test, Tukey-adjusted). The 50/10 and 10/10 treatment groups were not significantly different from one another on average (*p* > 0.05, Student’s *t*-test, Tukey-adjusted), further supporting the importance of nitrate concentration in determining phosphate uptake in plants. This data provides evidence for nitrate-phosphorous interactions in grasses that may be analogous to what has been described in Arabidopsis. Additionally, this data indicates that there are likely important effects of abiotic stress on root phenotypes that warrant future research.

**Figure 6.**
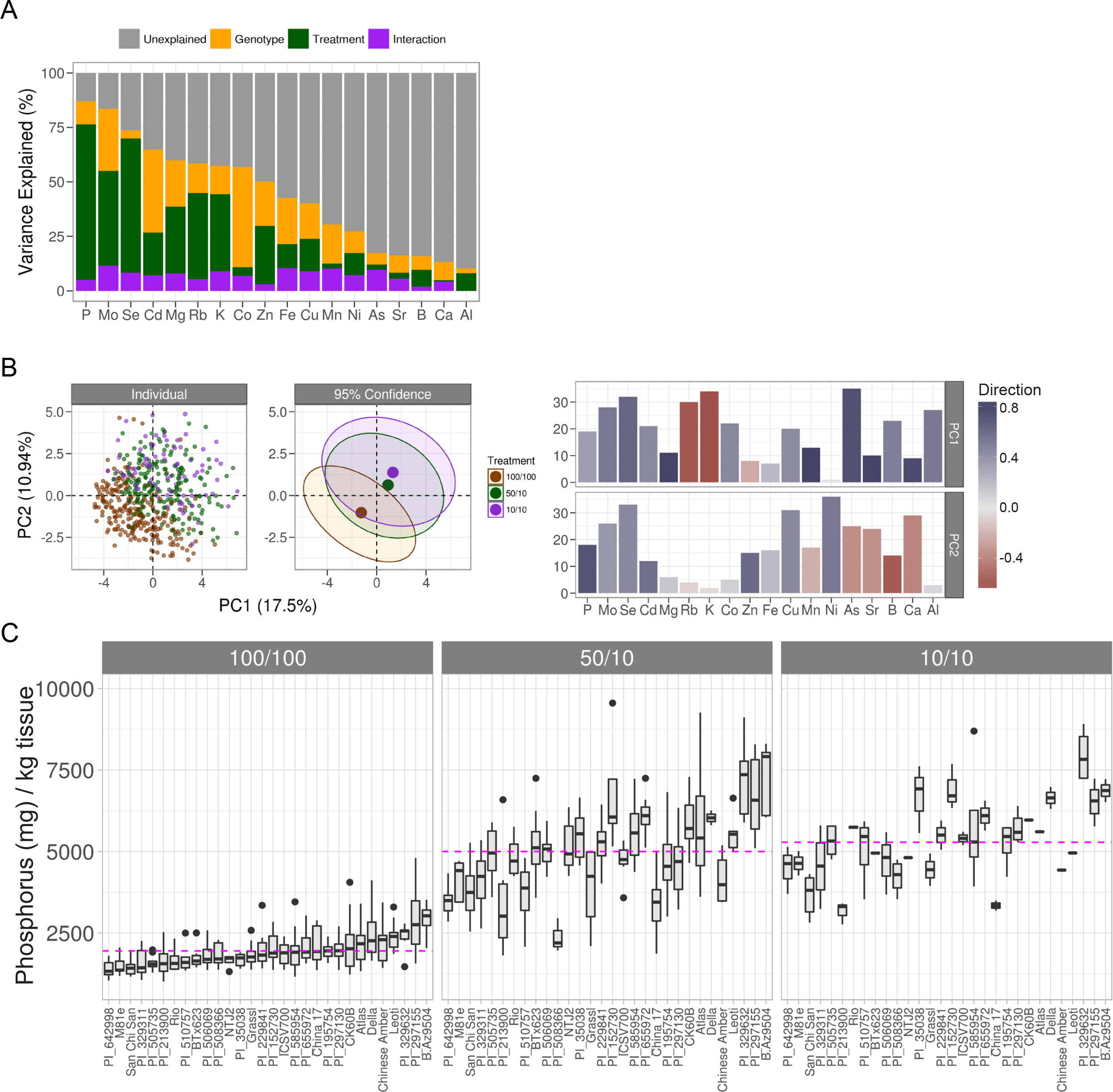
Ionomic profiling of genotypes at the end of the experiment. A) The percent variance explained by each partition of the total variance model (above). B) Left: PCA plots (all elements) colored by treatment for individual genotypes (left) and 95% confidence ellipses (right). The percent variance explained by each component is indicated in parentheses. Right: Loadings for each element from the first two PCs are shown on the y-axis and are color filled based on the direction and strength of the contribution. Positive direction is colored blue and negative direction is colored red. For a given element, the color for PC1 and PC2 are related by the unit circle and saturation of the color is equal to the length of the projection into each of the two directions. C) Boxplots representing dry weight concentrations for all elements and all nitrogen treatments. Concentrations are reported as parts-per-million (y-axis: mg analyte/kg sample) for each genotype (x-axis). The median is indicated by a black bar within each box. Magenta line: mean phosphorous concentration for given treatment group.

## DISCUSSION

Crops adapted to nutrient-poor conditions will be an invaluable resource for realizing the goal of dedicated bioenergy crops grown without irrigation and limited fertilizer on marginal lands. Robust, quantitative phenotypes are a prerequisite for genetic investigations and these can be gathered using high throughput phenotyping and image analysis. In order to test for and quantify G × E interactions we designed a strategy that utilized tightly controlled environmental conditions in a high-throughput manner in the genetically diverse, stress-tolerant crop, sorghum. We characterized changes in plant size and color over time as well as elemental profile as outputs of stress tolerance. Importantly, this work is intended to not only produce insights into sorghum biology and crop improvement, but also serve as a resource and an important step forward for high-throughput phenotyping in plants, providing analysis tools to the community as a whole.

One important question that remains is how plants efficiently utilize available nitrogen. Previous work has shown that plants use different forms of nitrogen, yet preference can be influenced greatly by genotype and the environment. Factors such as soil pH, CO_2_ levels, temperature and the availability of other nutrients have an impact on nitrogen uptake (Jackson and Reynolds, 1996; Coskun et al., 2016). Additionally, root architecture is affected by nitrogen source and nutrient availability. It has been shown for a number of species, including maize and barley, that ammonium causes a reduction in lateral root branching that can be reversed with the addition of phosphorous (Drew, 1975; Ma et al., 2013; Thomas et al., 2016; Giles et al., 2017). Compounding this equation, ammonium also causes acidification of the soil, which affects the uptake of other nutrients and likely alters the root microbiome, further complicating most analysis. Under the tested experimental conditions, some genotypes were more affected by nitrogen source in terms of end biomass than others, for example the difference between San Chi San and Atlas (Figure 4C). Also to this point, we show that phosphorous was one of the elements with the largest treatment effect and that the measured concentrations of phosphorous were higher in the low nitrogen treatment groups, both of which received the same nitrate treatment, compared to the high nitrogen treatment group (Figure 6). Taken together, these data are consistent with what other studies that have shown: some genotypes have a preference for nitrogen source and other environmental factors influence that preference. The interdependence between nitrogen uptake and phenotypic output in plants highlights the necessity of high-throughput, tightly controlled studies for answering these and other fundamental questions.

Some of the most productive crops in use today are C4 grasses like corn (*Zea mays*), sorghum (*Sorghum bicolor*), and sugarcane (primarily *Saccharum officinarum*) (Reviewed in Leakey, 2009). These crops have cellular functions and chemistries that result in high rates of photosynthesis in spite of drought and nutrient-poor conditions. However, within each crop group, significant genetic and phenotypic variety exists. The sorghum diversity panel presented here represents a wide, yet incomplete, range of known sorghum genotypic and phenotypic diversity. Tens of thousands of sorghum accessions are curated and maintained by a number of national and international institutions (Kimber et al., 2013). The largest such institution, the US National Sorghum Collection (GRIN database), provides agronomic characteristic information for 40–60% of the collection (e. g. growth and morphology characteristics, insect and disease resistance, chemical properties, production quality, photoperiod in temperate climates). Thus, much work is yet to be done to fully characterize and maximize the potential of this hearty, productive crop species.

Nitrogen use efficiency is traditionally defined by the difference in biomass or grain production between plants grown in resource sufficient versus resource limited conditions at the end of the growing season. Stated differently, this measure asks the question: How efficient is a plant at translating a provided resource (nitrogen) into plant biomass. Equally important is the ability to efficiently use a limited resource. Factors that play into these distinct definitions of resource use efficiency include ability to survive periods of extreme stress and rapid utilization of resources as they become available. In this manuscript, we make progress toward deconstructing the building blocks that make up nitrogen response phenotypes. These analyses reveal diverse quantitative indicators of abiotic stress and genotypic differences in stress mitigation that can be used to further crop improvement. Having made progress toward deconstructing these building blocks, we are now in a position to discover the underlying genetic explanations for genotypic variability in resource use efficiency and tolerance to resource limited growth conditions. This work forms a foundation for future research to overlay additional abiotic and biotic stress conditions to achieve a holistic view of sorghum G × E phenotypes. The overall goal of this research is to support such efforts and expedite the process of meaningful crop improvement.

## CONCLUSION

Plant stress tolerance is important for food security and sorghum has potential as a high-yielding, stress-tolerant crop. ‘Resource use efficiency’ is often measured in one of two ways: 1) a comparison between yield production under resource-sufficient and resource-limited conditions, 2) the ability to survive within resource limited environments. Here we describe and apply high-throughput phenotyping methods and element profiling to sorghum grown under variable nutrient levels. We quantify nitrogen use efficiency in genetically diverse sorghum accessions based on color fluctuations and growth rate over time and elemental profile. Through this analysis we report a time-efficient, robust approach to identifying resource use efficient and abiotic stress tolerant plants.

## MATERIALS AND METHODS

### Plant growth conditions

Round pots (10 cm diameter) fitted with drainage trays were pre-filled with Profile^®^ Field & Fairway™ calcined clay mixture (Hummert International, Earth City, Missouri) the goal being to minimize soil contaminates (microbes, nutrients, etc.) and maximize drainage. Before the beginning of the experiment, the thirty genotypes of *Sorghum bicolor* (L.) Moench (Table S1) were planted, bottom-watered once daily using distilled water (reverse osmosis), then allowed to germinate for 6 days in a Conviron growth chamber (day/night temperature: 32ºC/22ºC, day/night humidity: 40%/50% (night), day length: 16hr, light source: Philips T5 High Output fluorescent bulbs (4100 K (Cool white)) and halogen incandescent bulbs (2900K (Warm white)), light intensity: 400 µmol/m^2^/s). On day 6, plants were barcoded (including genotype identification, treatment group, and a unique pot identification number), randomized, then loaded onto the Bellwether Phenotyping Platform (Conviron, day/night temperature: 32ºC/22ºC, day/night humidity: 40%/50% (night), day length: 16hr, light source: metal halide and high pressure sodium, light intensity: 400 µmol/m^2^/s). Plants continued to be watered using distilled water by the system for another 2 days, with experimental treatments (described below) and imaging beginning on day 8.

### Nitrogen treatments

100/100 (100% Ammonium/100% Nitrate): 6.5 mM KNO_3_, 4.0 mM Ca(NO_3_)_2_·4H_2_O, 1.0 mM NH_4_H_2_PO_4_, 2.0 mM MgSO_4_·7H_2_O, micronutrients, pH ~4.6

50/10 (50% Ammonium/10% Nitrate): 0.65 mM KNO_3_, 4.95 mM KCl, 0.4 mM Ca(NO_3_)_2_·4H_2_O, 3.6 mM CaCl_2_·2H_2_O, 0.5 mM NH_4_H_2_PO_4_, 0.5 mM KH_2_PO_4_, 2.0 mM MgSO_4_·7H_2_O, micronutrients, pH ~4.8

10/10 (10% Ammonium/10% Nitrate): 0.65 mM KNO_3_, 4.95 mM KCl, 0.4 mM Ca(NO_3_)_2_·4H_2_O, 3.6 mM CaCl_2_·2H_2_O, 0.1 mM NH_4_H_2_PO_4_, 0.9 mM KH_2_PO_4_, 2.0 mM MgSO_4_·7H_2_O, micronutrients, pH ~5.0

### The same micronutrients were used for all above treatments

4.6 µM H_3_BO_3_, 0.5 µM MnCl_2_·4H_2_O, 0.2 µM ZnSO_4_·7H_2_O, 0.1 µM (NH_4_)_6_Mo_7_O_24_·4H_2_O, 0.2 µM MnSO_4_·H_2_O, 71.4 µM Fe-EDTA

### Image Processing

Images were analyzed by using an open-source platform named PlantCV ((Fahlgren et al., 2015), http://plantcv.danforthcenter.org). This package primarily contains wrapper functions around the commonly used open-source image analysis software called OpenCV (version 2.4.5). To get useful information from a given image, the plant must be segmented out of the picture using various mask generation methods to remove the background so all that remains is plant material (see Figure 1). A pipeline was developed to complete this task for the side-view and top-view cameras separately and they were simply repeated for every respective image in a high-throughput computation cluster. For this dataset of approximately 90,000 images with the computation split over 40 cores, computation time was roughly four hours. Upon completion, data files are created that contain parameterizations of various shape features and color information from several color-spaces for every image analyzed.

### Outlier Detection and Removal Criteria

Each treatment group began with 9 reps per genotype for the 100/100 and 50/10 treatment groups and 6 reps per genotype for the 10/10 treatment group. Outliers were detected and removed by implementing Cook’s distance on a linear model (Cook, 1977) that only included the interaction effect of treatment, genotype and time. That is, for each observation (every image, for every plant, every day), an influence measure is obtained as the difference of the model with and without the observation. After getting a measure for all observations in the dataset, outliers were defined as having an influence greater than four times that of the mean influence and were subsequently removed from the remaining analysis. In total 5.8% of the data, 1598 images, was removed using this method.

### PCA

Three types of PCA’s are generated: one for the shape features, color features, and ionomics. All shape parameterizations that are generated from PlantCV are included in the dimensional reduction. Principle components of color, as defined by the hue channel in two degree increments, is examined using all one hundred eighty bins in the dimensional reduction. Ionomics PCA was generated using every element that passed internal standards of quality.

### GLMM-ANOVA

Using area as the response variable, a general linear mixed model was created to identify significance sources of variance adjusting for all other sources, otherwise known as type III sum of squares. Designating genotype as G, treatment as E, and time as T, there are six fixed effects: G, E, GxE, GxT, ExT, GxExT. The mixed effect is a random slope and intercept of the repeated measures over time. Wald Chi-Square statistic was implemented and is a leave-one-out model fitting procedure which allows for adjustment of all other sources.

### Heatmaps

Every cell is a comparison of treatments using a 1-way ANOVA wherein the *p*-value is obtained from a F-statistic generated from the sum of squares of the treatment source of variation. After getting all the raw *p*-values, a Benjamini-Hochberg FDR multiple comparisons correction is done to aid in eliminating false positives. The *p*-value distribution was very left skewed so a log-transform is used to normalize them. Agglomerative, hierarchical clustering was used on the corrected *p*-values. Each genotype had an associated vector of *p*-values and a Canberra distance is calculated for all pairwise vectors which are then grouped by Ward’s minimum variance method.

### Color Processing

PlantCV returns several color-space histograms for every image that is run through the pipeline (RGB, HSV, LAB, and NIR). Every channel from each color-space is a vector representing values (or bins) from 0 to 255 which are black to full color respectively. All image channel histograms were normalized by dividing each of the bins by the total number of pixels in the image mask ultimately returning the percentage of pixels in the mask that take on the value of that bin. The hue channel is a 360 degree parameterization of the visible light spectrum and contains the number of pixels found at each degree. The colors of most importance are between 0 and 120 degrees which correspond to the gradient of reds to oranges to yellows to greens. Colors beyond this range, like cyan and magenta, have values of all zeros and are not shown. Means and 95% confidence intervals as calculated on a per degree basis over the replicates. Area under the curve calculations were done using the trapezoidal rule within the two ranges of 0 to 60 degrees and 61 to 120 degrees which are designated as yellow and green peaks respectively.

### Ionomics Profiling and Analysis

The most recent mature leaf was sampled from each plant on day 26 of each experiment, placed in a coin envelope and dried in a 45ºC oven for a minimum of 48 hours. Large samples were crushed by hand and subsampled to 75mg. Subsamples or whole leaves of smaller samples were weighed into borosilicate glass test tubes and digested in 2.5 mL nitric acid (AR select, Macron) containing 20ppb indium as a sample preparation internal standard. Digestion was carried out by soaking overnight at room temperature and then heating to 95ºC for 4hrs. After cooling, samples were diluted to 10 mL using ultra-pure water (UPW, Millipore Milli-Q). Samples were diluted an additional 5x with UPW containing yttrium as an instrument internal standard using an ESI prepFAST autodilution system (Elemental Scientific). A Perkin Elmer NexION 350D with helium mode enabled for improved removal of spectral interferences was used to measure concentrations of B, Na, Mg, Al, P, S K, Ca, Mn, Fe, Co, Ni, Cu, Zn, As, Se, Rb, Mo, and Cd. Instrument reported concentrations are corrected for the yttrium and indium internal standards and a matrix matched control (pooled leaf digestate) as described (Ziegler et al., 2013). The control was run every 10 samples to correct for element-specific instrument drift. Concentrations were converted to parts-per-million (mg analyte/kg sample) by dividing instrument reported concentrations by the sample weight.

Outliers were identified by analyzing the variance of the replicate measurements for each line in a treatment group and excluding a measurement from further analysis if the median absolute deviation (MAD) was greater than 6.2 (Davies and Gather, 1993). A fully random effect model is created for every element and partial correlations are calculated for treatment, genotype and the interaction using type-III sum of squares.

## ACKNOWLEDGEMENTS

We acknowledge Mindy Darnell and Leonardo Chavez from The Bellwether Foundation Phenotyping core facility at the Danforth center as well as Diana Fasanello and Molly Kuhs for their assistance in running the experiments. We would also like to thank Dr. Greg Ziegler for his help with the ionomics analysis and Dr. Stephen Kresovich for his many helpful discussions and for supplying the seed for the sorghum diversity panel.

## SUPPLEMENTAL DATA

**Tab. S1.** Genotypic information for accessions included in this study.

**Fig. S1.** All shape parameterizations returned from PlantCV had correlations calculated to all other shapes. Correlation is on a scale from -1 to 1 indicating inversely or directly correlated and is being shown in color from red to blue. Radius of the circle in each cell is on a scale between 0 and 1 which corresponds to the absolute value of the correlation.

**Fig. S2.** Tables showing results of ANOVA indicating significance of experimental variation explained by either genotype, type, photoperiod or race as found by Wald’s Chi-Square tests with their associated degrees of freedom (DF). Significant *p*-value < 0.1, bold. All three nitrogen treatments are included in the calculations.

**Fig. S3.** Statistical analysis of differences between 50/10 and 10/10 groups from the nitrogen deprivation experiment in area over time (bottom, plant age) for the 30 sorghum genotypes analyzed. *q*-values for the heat map are indicated in blue, with darkest coloring representing most significance. The Canberra distance-based cluster dendrogram (right) was generated from calculated *q*-values.

**Fig. S4.** Color changes in individual late and early responding genotypes when the peak experimental effects were observed (day 13). To make the figure average histograms from the indicated genotypes within the 100% and 10% treatment groups were subtracted from one another. Grey areas indicate standard error.

**Fig. S5.** Boxplots representing dry weight concentrations for all elements and all nitrogen deprivation treatments. Concentrations are reported as parts-per-million (y-axis: mg analyte/kg sample) for each genotype (x-axis).

## Supplemental files

Processed data: sorg_nitrogen_all_shapes.csv – Processed shape data from nitrogen experiment. Ionomics_RawData_Nitrogen.csv – Processed ionomics data from nitrogen experiment.

## Notes

Funding: This work was supported by US Department of Energy award DE-SC0014395.

